# Spindle pole proteins confine chromosomes to ensure their expulsion during female meiosis

**DOI:** 10.64898/2026.03.24.713942

**Authors:** Kan Yaguchi, Lifeng Chen, Abdollah Mohammadi-Sangcheshmeh, Judith Tafur, Nicole E. Familiari, Edward J. Grow, Michael K. Rosen, Jeffrey B. Woodruff

**Affiliations:** Dept. of Cell Biology, UT Southwestern Medical Center, Dallas, TX 75390, USA; Dept. of Biophysics, UT Southwestern Medical Center, Dallas, TX 75390, USA; Cecil H. and Ida Green Center for Reproductive Biology Science, UT Southwestern Medical Center, Dallas, TX 75323, USA; Dept. of Obstetrics and Gynecology, UT Southwestern Medical Center, Dallas, TX 75323, USA; Howard Hughes Medical Institute, UT Southwestern Medical Center, Dallas, TX 75390, USA

## Abstract

Animal oocytes undergo highly asymmetric divisions to expel excess copies of their genome into compact cells called polar bodies. This requires tight clustering and cortical positioning of meiotic chromosomes, yet the mechanism remains incompletely understood. Using *C. elegans* oocytes, we found that the meiotic spindle pole protein ZYG-9/ch-TOG delocalizes from microtubules to spread along the surface of chromosomes and prevent their dispersal in meiotic anaphase. This effect was more pronounced in the absence of the spindle, where ZYG-9 formed into a micron-scale droplet that enveloped all chromosomes. Purified ZYG-9 was sufficient to bind DNA and coat reconstituted chromatin. Mutations that perturb ZYG-9-DNA binding impaired chromosome packaging into polar bodies, resulting in oocytes carrying extra chromosomes and reduced fertility. We propose that liquid-like assemblies of spindle pole proteins are repurposed as surface-acting glue to tightly package meiotic chromosomes into polar bodies, thus ensuring oocytes have the correct genome copy number.

## Introduction

Female meiosis represents one of the most extreme cases of asymmetric cell division in animals. In this process, an oocyte divides two times without an intervening gap phase, thus creating one large haploid egg and two compact cells called polar bodies. Such division reduces the number of chromosomes in half while retaining maternally supplied cytoplasm needed for rapid mitotic divisions after fertilization of the egg. Polar bodies can be 1000-2000X smaller in volume than the egg in many species (1–4). Polar body size negatively correlates with egg viability in patients undergoing *in vitro* fertilization (5, 6). Thus, creating a compact polar body during meiosis is an important and conserved mechanism to ensure animal fertility. This process requires clustering meiotic chromosomes near the oocyte periphery and tightly packaging them into the limited space of the polar body. The mechanisms that spatially confine chromosomes in the vast oocyte cytoplasm and ensure their retention in the polar bodies are incompletely understood.

The microtubule-rich spindle has been considered the canonical spatial constraint for meiotic chromosomes. After germinal vesicle breakdown, microtubule polymerases and regulators enrich around meiotic chromosomes to nucleate microtubules (7–10). During prometaphase and metaphase, the microtubules first form a cage around chromosomes and subsequently organize into an acentrosomal bipolar spindle (7, 11). However, even when the spindle microtubules are artificially depolymerized during metaphase, individual meiotic chromosomes remain in spatial proximity (12–14). This suggests the existence of a microtubule-independent spatial constraint for meiotic chromosomes. During the transition from metaphase to anaphase, microtubules disappear at the spindle poles and then reappear between sister chromosomes, coincident with a decrease in the overall spindle size and tighter clustering of chromosomes (15–17). The restructured spindle then exerts a pushing force to segregate chromosomes towards the pole region and into the forming polar body (16, 17). Based on these observations, we hypothesized that a spindle pole-associated factor might direct tight clustering of chromosomes within oocytes.

## Results

### The microtubule polymerase ZYG-9 associates with anaphase chromosomes in *C. elegans* oocytes

A core set of conserved microtubule-associated proteins concentrate at the meiotic spindle pole throughout meiosis (e.g., calmodulin/CMD-1, ASPM/ASPM-1, NuMA/LIN-5, Tacc3/TAC-1 and chTOG/ZYG-9) (7, 11, 18) (Figure S1A). Some of these proteins assemble into dynamic droplets, which wet the microtubules and eventually extend from the microtubule lattice to fuse with other droplets (18). It is unclear whether these spindle pole proteins associate with chromosomes during meiotic completion. Thus, we tracked endogenously fluorescent-tagged proteins in *ex utero C. elegans* oocytes using time-lapse imaging and super-resolution microscopy (Figure 1A, 1D; Figure S1A). During metaphase, the spindle poles were spatially separated from the chromosomes (Figure 1A) (19). At the anaphase transition, indicated by spindle rotation and shortening, spindle pole proteins accumulated and spread along the chromosome surface (Figure 1A-C). This was also apparent when visualized from an orthogonal orientation (Figure S1B). These spindle pole proteins remained on the chromosome surface during late anaphase in meiosis I, as indicated by fluorescent line scans (Figure 1D). These results suggest a secondary role for the spindle pole beyond their canonical microtubule-organizing functions during meiotic anaphase.

**Figure 1.**
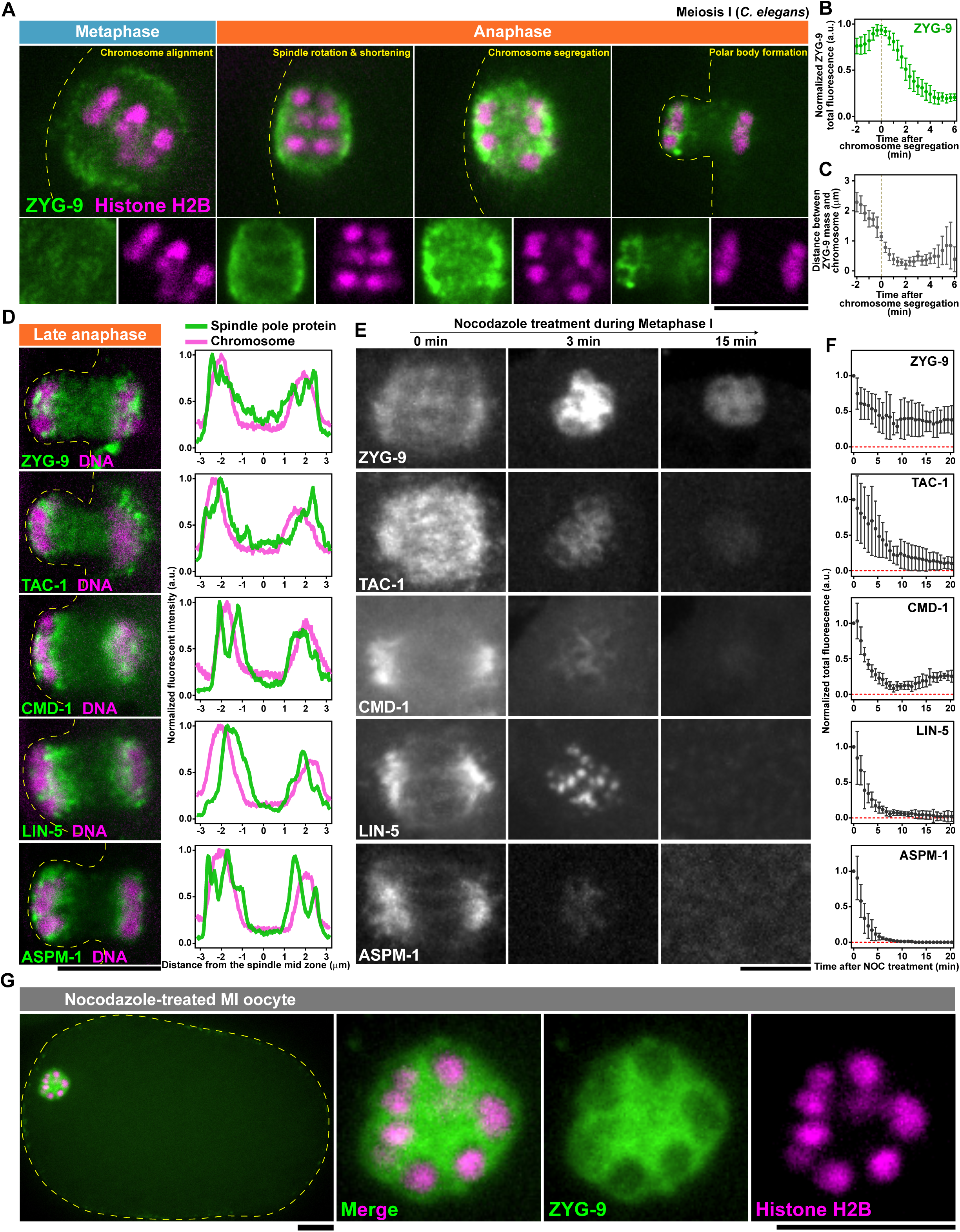
ZYG-9 coats meiotic chromosomes during anaphase in *C. elegans* oocytes. **(A)** Super-resolution microscopy of *ex utero C. elegans* oocytes expressing GFP::ZYG-9 (spindle poles, green) and mCherry::H2B (chromosomes, magenta) in meiosis I (top). Chromosome regions are cropped (bottom). **(B)** Total fluorescence of spindle-associated GFP::ZYG-9. **(C)** Distance between the spindle pole and the chromosome bivalent. Data are normalized to maxima and represent mean ± 95% C.I. (n= 10). **(D)** Super-resolution microscopy of endogenously labeled spindle pole proteins (green) and chromosomes (Sir-DNA, magenta) during chromosome segregation in anaphase (left). Line scans of normalized signal intensity across the long axis of the spindle (right). **(E)** Time-lapse imaging of spindle pole proteins after nocodazole treatment. **(F)** Normalized total fluorescence of each spindle pole proteins in (E). Data represent mean ± 95% C.I. of GFP::ASPM-1 (n= 6), CMD-1::GFP (n= 8), LIN-5::mNeonGreen (n= 6), GFP::TAC-1 (n= 5), GFP::ZYG-9 (n= 5). **(G)** Super-resolution microscopy of *ex utero* oocytes treated with 20 µM nocodazole for 15 min (left). Enlarged images shown on the right. All scale bars are 5 µm.

Previous room temperature electron microscopy showed that microtubules are largely absent from the spindle pole and instead prominently appear between chromosomes during meiotic anaphase (16, 17). It is therefore possible that spindle pole proteins localize on anaphase chromosomes independently of microtubules. To test this hypothesis, we depolymerized microtubules just before the metaphase-anaphase transition using nocodazole and visualized spindle pole proteins (Figure 1E). While ASPM-1, CMD-1, LIN-5, and TAC-1 instantly dispersed after nocodazole addition, ∼35% of the ZYG-9 pool that originally localized within the spindle reformed into a micron-scale mass (Figure 1E, 1F; Video S1). This ZYG-9 mass had multiple internal voids occupied by chromosomes (Figure 1G). Thus, ZYG-9 is unique among spindle pole proteins in that it superficially associates with chromosomes both in the presence and absence of the meiotic spindle in *C. elegans* oocytes.

We next examined the dynamics of the ZYG-9 assembly around chromosomes. The ZYG-9 mass photo-recovered after partial and full bleaching, indicating internal and external molecular exchange (Figure S2A and S2B). Upon nocodazole treatment, ZYG-9 sometimes appeared as separate droplets that coalesced into a larger spherical drop (Figure S2C). ZYG-9 droplets also wetted and bent the plasma membrane upon the contact (Figure S2C). These behaviors are classical features of biomolecular condensates (20, 21). These data revealed that, when not constrained by spindle microtubules, ZYG-9 acts as a micron-scale, liquid-like condensate that encapsulates all meiotic chromosomes.

### ZYG-9 acts as a chromosome-trapping glue during meiotic anaphase I

Depletion or inactivation of ZYG-9 in metaphase or prior was reported to cause spindle pole destabilization and the blockage of meiotic progression, indicating that ZYG-9 organizes microtubules to help form and maintain a functional bipolar spindle (22, 23). However, the importance of ZYG-9 specifically during the anaphase transition and beyond, when ZYG-9 coats chromosomes, was not tested. To test if ZYG-9 plays a functional role in chromosome maintenance, we used time-controlled degradation of ZYG-9 tagged with an auxin-inducible degron (degron::GFP::ZYG-9, Figure 2A). When we added auxin in early metaphase, prior to spindle rotation, ZYG-9 was degraded in ∼4 min yet chromosomes remained close to each other for over 20 min (Figure 2A-D). In contrast, when we added auxin during the anaphase transition (i.e., after spindle rotation), chromosomes dispersed and drifted away from each other in the large oocyte cytoplasm (Figure 2A-D). The severity of this phenotype scaled in relation to the timing of auxin addition: waiting to degrade ZYG-9 until the spindle fully rotated led to the most dramatic chromosome dispersal (Figure 2E). Thus, ZYG-9 is required to spatially constrain chromosomes when the meiotic spindle rotates and reorganizes during the anaphase transition.

**Figure 2.**
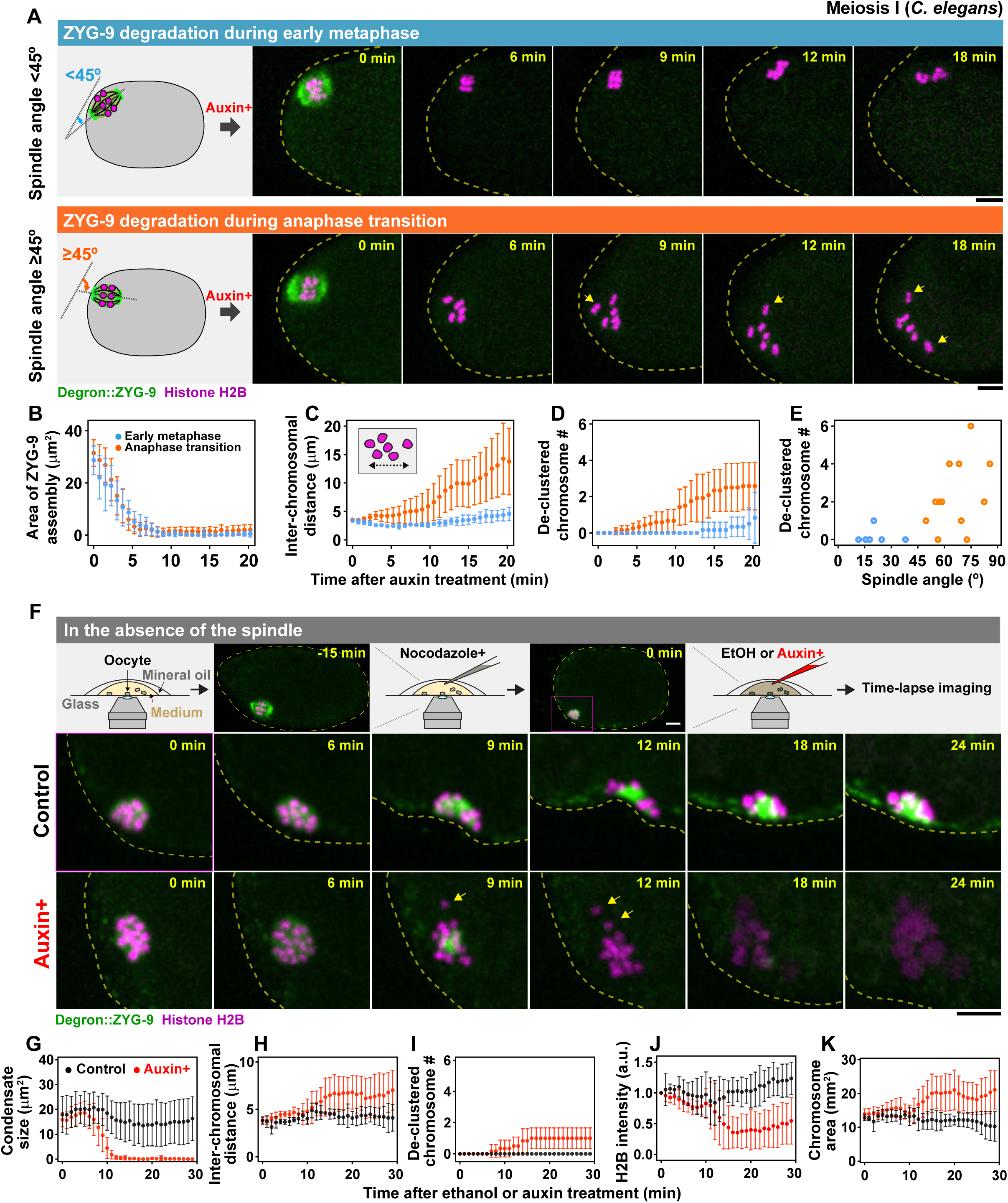
ZYG-9 acts as chromosome-trapping glue during meiosis. **(A)** Auxin-induced degradation of degron::GFP::ZYG-9 before (top) and during (bottom) spindle rotation at the metaphase-anaphase transition. Data were sorted by the angle of the long axis of the spindle against the tangent line of the oocyte membrane (left schemes). **(B-E)** Quantification of spindle-associated degron::GFP::ZYG-9 disassembly (B), distance between the farthest chromosomes (C), and number of de-clustered chromosomes (D) after auxin-treatment. Data represent mean ± 95% C.I. (before spindle rotation, blue dots, n= 6; during spindle rotation, orange dots, n= 12). The number of de-clustered chromosomes 15 min after auxin-treatment was plotted against the spindle angle (E). **(F)** Diagram of the experimental procedure to disrupt ZYG-9 condensates (top). Oocytes were pre-treated with nocodazole for 15 min, followed by the ethanol (middle) or auxin treatment (bottom). **(G-K)** Quantification of ZYG-9 condensate size (G), distance between the farthest chromosomes (H), number of de-clustered chromosomes (I), normalized mCherry::H2B intensity (J), and chromosome size (K) after ethanol or auxin treatment. Data represent mean ± 95% C.I. (Ethanol, black dots, n= 5; auxin, red dots, n= 7). All scale bars are 5 µm. Yellow arrows indicate de-clustered chromosomes.

To test if this chromosome organization stems from the microtubule polymerase activity of ZYG-9, we acutely depleted ZYG-9 in nocodazole-treated metaphase I oocytes, where chromosomes are encapsulated by a ZYG-9 condensate in the spindle-less oocyte (Figure 2F). In control oocytes (no auxin, 0.5% ethanol), the ZYG-9 condensate stably tethered all meiotic chromosomes (Figure 2F-I). In contrast, when the condensates disappeared after auxin-induced ZYG-9 degradation (Figure 2F-G), chromosomes were modestly dispersed (Figure 2H and 2I). We conclude that the tethering activity of ZYG-9 does not require microtubules. Unexpectedly, chromosomes underwent decompaction after ZYG-9 degradation, as indicated by an increase in size and a decrease in histone intensity (Figure 2J and 2K). This differs from ZYG-9 degradation in the presence of microtubules (Figure 2A). Thus, the spindle and ZYG-9 act in parallel to maintain chromosome compaction. Taken together, our data suggest that liquid-like assemblies of ZYG-9 associate with and spatially confine meiotic chromosomes independent of microtubules.

### ZYG-9 directly binds DNA to form co-condensates *in vitro*

We sought to investigate how ZYG-9 associates with meiotic chromosomes. The most obvious potential docking site is the outer kinetochore, which is loaded around the entirety of *C. elegans* chromosomes (24, 25). Homologs of ZYG-9 were reported to bind the outer kinetochore-localized Ndc80 in budding yeast and human somatic cells (26–29). However, ZYG-9 still accumulated around chromosomes in *ndc-80(RNAi)* oocytes—where the outer kinetochore is lost (30)—upon nocodazole treatment (Figure 3A and 3B). Thus, ZYG-9 assembles on the chromosome surface independently of the outer kinetochore.

**Figure 3.**
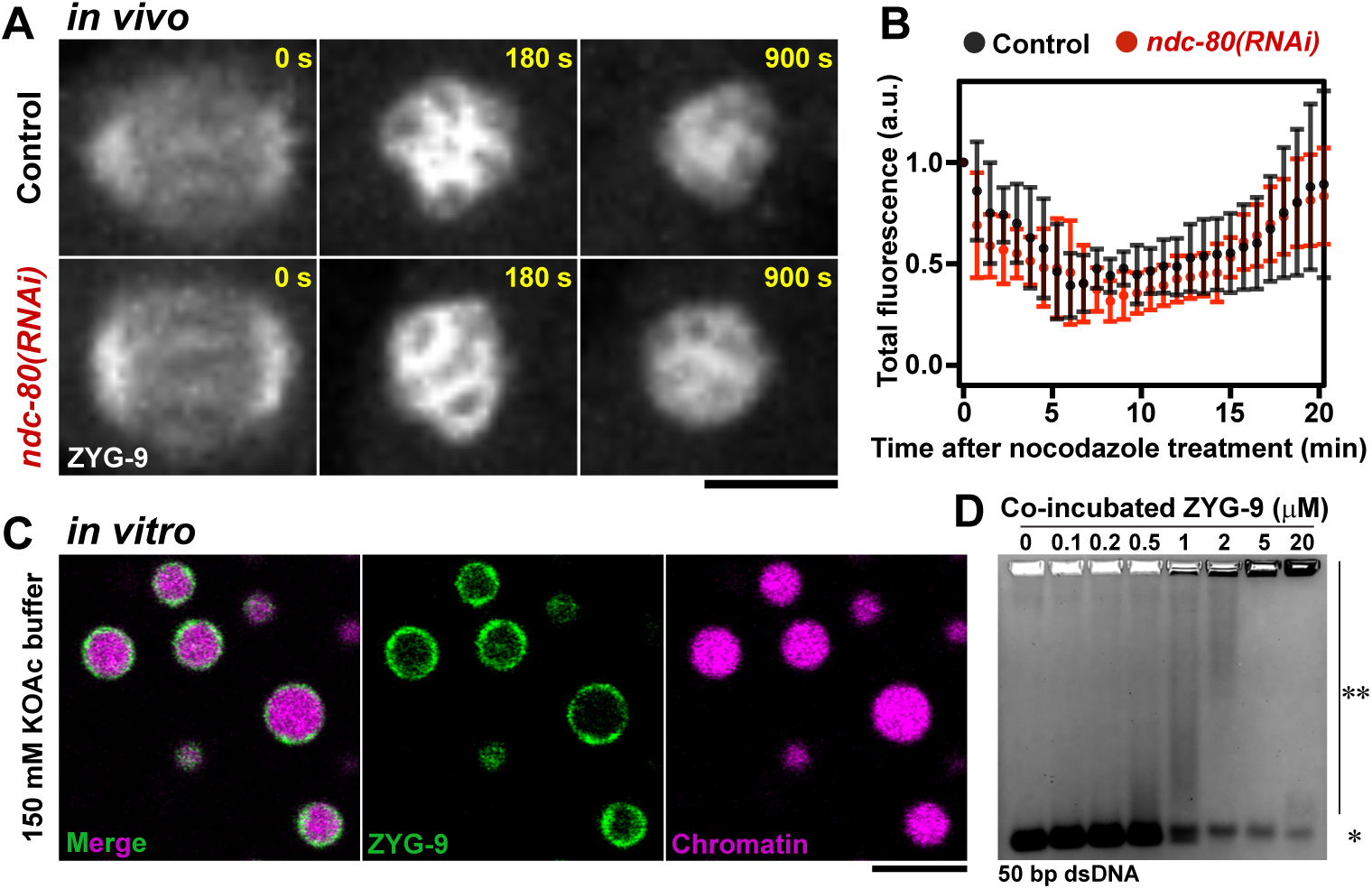
ZYG-9 is sufficient to bind DNA and encapsulate condensed chromatin *in vitro*. **(A)** Control and *ndc-80(RNAi)* oocytes expressing GFP::ZYG-9 were treated with nocodazole. **(B)** Total fluorescence of chromosome- and spindle-localized GFP::ZYG-9 in (A). Data represent mean ± 95% C.I. (Control, n= 9; *ndc-80(RNAi)*, n= 8). **(C)** *In vitro* reconstitution of purified GFP::ZYG-9 proteins and chromatin under near physiological salt concentration (150 mM KOAc). 500 nM protein and 500 nM chromatin were co-incubated for 30 min. **(D)** Electrophoretic mobility shift assay of 0.2 µM 6-FAM-labeled 50 bp dsDNA co-incubated with 0-20 µM ZYG-9. Unbound (*) and bound (**) species are indicated. All scale bars are 5 µm.

Alternatively, ZYG-9 may directly bind the chromosomes independently of other meiotic components. To test this hypothesis, we reconstituted dodecameric nucleosome arrays using purified histone octamers and linear dsDNA with 12 Widom’s 601 sequence (31, 32). This synthetic chromatin spontaneously condenses and assembles into dense micron-scale droplets under near physiological salt concentrations, recapitulating the supramolecular architecture of *in vivo* chromosomes (33, 34). When purified GFP::ZYG-9 was incubated with chromatin, ZYG-9 coated the surface of the chromatin droplets (Figure 3C, S3A). This result suggests that self-associating chromatin excludes ZYG-9 from the interior yet interacts with ZYG-9 on the surface. The superficial ZYG-9 phase was dynamic and rearranged upon contact with other droplets to allow fusion of interior chromatin (Video S2). Thus, purified nucleosome arrays and ZYG-9 proteins form a two-phase system that mimics ZYG-9 encapsulation of chromosomes seen *in vivo* (Figure 1G).

As an orthogonal approach to test if ZYG-9 directly binds DNA, we performed an electrophoretic mobility-shift assay (EMSA). Addition of pure ZYG-9 slowed the migration of 50-bp polyA dsDNA in a concentration-dependent manner, indicating protein-DNA complex formation (Figure 3D). ZYG-9 also bound 5-kbp linearized dsDNA (Figure S3B). These results indicate that ZYG-9 is a novel DNA binding protein.

Incubation of DNA with higher concentrations of ZYG-9 induced formation of large complexes that were retained in the loading well of the gels (Figure 3D and S3B). Mass photometry revealed that, at 20 nM concentration, ZYG-9 formed primarily monomers and dimers (Figure S3C). Furthermore, using confocal imaging, we did not detect any micron-scale ZYG-9 assemblies at physiological protein concentration (500 nM) and salt concentration (150 mM KCl; Figure S3D). Adding 1 kbp dsDNA led to formation of robust, micron-scale co-condensates (Figure S3E and S3F). On the other hand, adding known ZYG-9-binding proteins (a/b tubulin, SPD-5, TAC-1) did not promote larger-scale assembly (Figure S3A). Photo-bleach experiments showed modest recovery of ZYG-9 proteins inside the co-condensates (Figure S3G). These results suggest that interactions with DNA promote higher-order assembly of ZYG-9 into viscous droplets, which has not been reported previously.

### An arginine patch within an unstructured linker of ZYG-9 is required for DNA binding and trapping *in vitro*

Next, we sought to define the DNA-binding motif of ZYG-9. ZYG-9 is a large multifunctional protein consisting of four helix-rich TOG family domains and a C-terminal helix, all of which are followed by intrinsically disordered regions (IDRs; Figure 4A). We first split ZYG-9 into 5 fragments, each containing a separate helix-rich domain and subsequent IDR (Figure 4A). We then purified these fragments and assessed their ability to bind 50 bp dsDNA by EMSA. Fragments #2 (a.a. 281-598, TOG2+IDR), #3 (a.a. 599-975, TOG3+IDR), and #5 (a.a. 1241-1415, C-terminal helix+IDR) bound DNA, albeit with different efficacy (Figure 4B). 20 µM of fragments #2 and #5 were required to detect a DNA-protein complex, whereas only 0.5 µM of fragment #3 was required (K_d_^app^ = 2.72 µM; Figure 4C). These results suggest that distinct domains of ZYG-9 bind DNA with different affinities.

**Figure 4.**
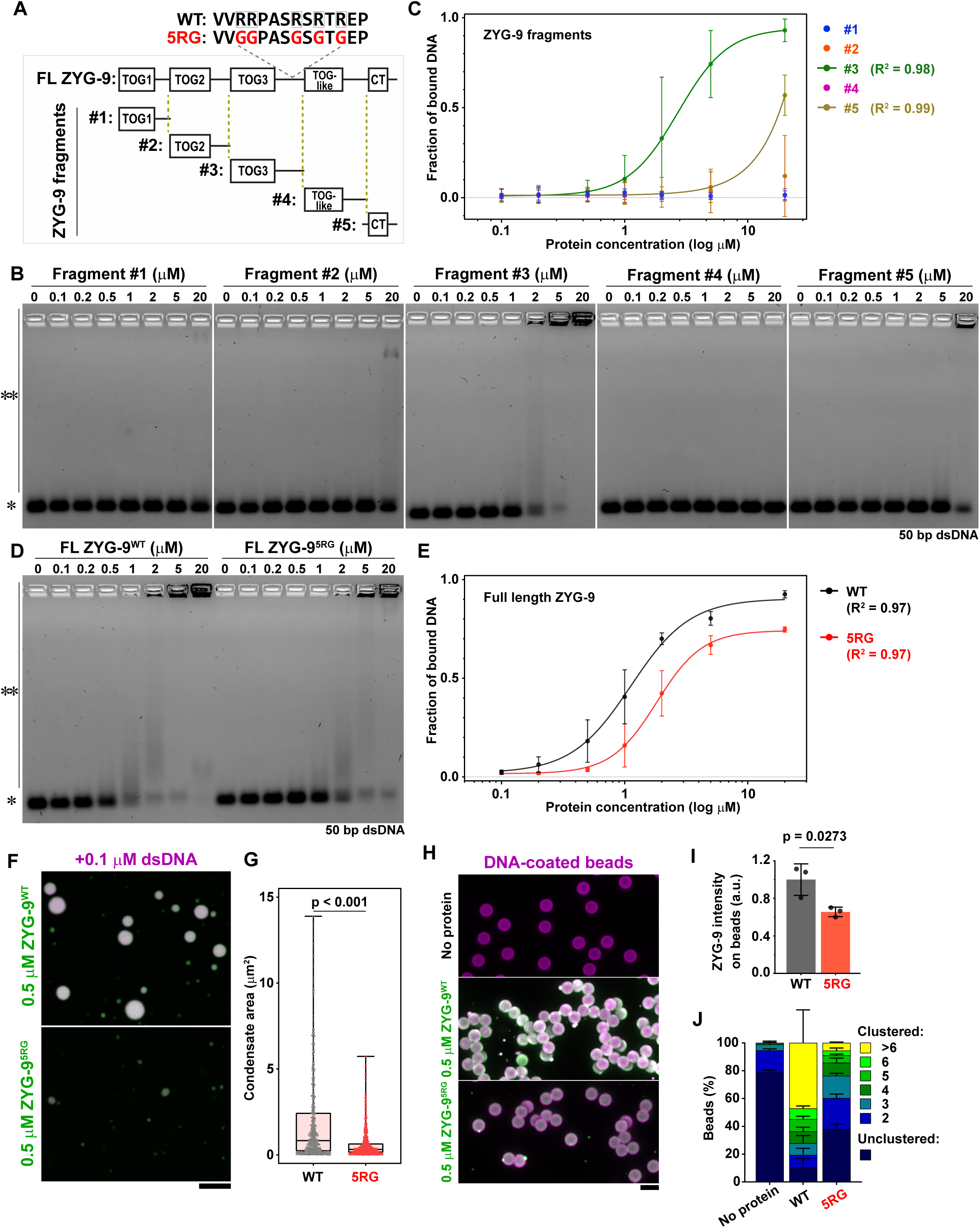
The arginine patch within ZYG-9’s IDR is needed for efficient DNA binding. **(A)** Diagram of purified ZYG-9 protein fragments (#1, a.a. 1-280; #2, a.a 281-598; #3, a.a. 599-975; #4, a.a. 976-1265; #5, a.a. 1241-1415) and a full-length ZYG-9 mutant (ZYG^5RG^: R895G, R896G, R900G, R902G, R904G). Helix-rich domains (boxes) and intrinsically disordered regions (IDRs, lines) are indicated. **(B-E)** Electrophoretic mobility shift assay of 0.2 µM 6-FAM-labeled 50 bp dsDNA co-incubated with 0-20 µM ZYG-9 fragments (B) or full-length proteins (D). Unbound (*) and bound (**) species are indicated. Bound fraction of dsDNA (C and E). Data represent mean ± 95% C.I. from at least three independent experiments and fitted curves with 4-parameter logistic model (R^2^ >0.97). **(F)** Confocal microscopy of full-length mScarlet::ZYG-9 proteins (green) co-incubated with Alexa 647-labeled 1 kbp dsDNA (magenta) *in vitro*. **(G)** Quantification of ZYG-9 condensate size in (F). Data represent condensate size with a box and whisker plot (WT, gray dots; 5RG, red dots). **(H-J)** Confocal microscopy of DNA beads in the absence (top) or presence of full-length mScarlet::ZYG-9 proteins (WT, middle; 5RG, bottom) *in vitro* (H). Intensity of superficial mScarlet::ZYG-9 on the beads (I) and proportion of clustered beads (J). Data represent mean ± SD from three independent experiments. p-value was determined by unpaired t-test. All scale bars are 5 µm.

To investigate the functional importance of ZYG-9-DNA binding, we aimed to mutate the DNA-binding motif without perturbing ZYG-9’s canonical spindle functions. TOG domains are required for microtubule polymerization (35, 36); the C-terminal helix and its tail serve as a docking site for the co-factor TAC-1 (37); and, the IDR following the TOG2 domain in human chTOG is required to regulate kinetochore-bound microtubules (38). Thus, of the three DNA-binding fragments we identified, only the IDR following TOG3 has no reported function. We found that this IDR contains a positively charged arginine patch (R895, R896, R900, R902, R904; Figure 4A); similar patches are found in other DNA-binding proteins (39, 40). Mutating these five arginines into glycines in the full-length protein (ZYG-9^5RG^) impaired efficient ZYG-9-DNA binding (Figure 4D and 4E; WT, K_d_^app^ = 1.12 µM; 5RG, K_d_^app^ = 1.83 µM). ZYG-9^5RG^ still formed co-condensates with 1 kbp DNA, but the droplet size was significantly decreased compared with those of ZYG-9^WT^ (Figure 4F and 4G). These results indicate that the arginine patch is required for ZYG-9-DNA binding, and that efficient ZYG-9-DNA interaction is required for robust co-condensation.

Our *in vivo* experiments indicated that perichromosomal ZYG-9 helps cluster meiotic chromosomes (Figure 2). To investigate whether ZYG-9 is sufficient for this activity, we co-incubated ZYG-9 with dsDNA-coated beads (Figure 4H), which do not coalesce spontaneously like the chromatin droplets (Video S2). ZYG-9^WT^ co-localized with 1 kbp DNA on the bead surface and promoted clustering (Figure 4H-J), indicating that superficial ZYG-9 can interact with ZYG-9 proteins on other beads to stick them together. On the other hand, for ZYG-9^5RG^, superficial accumulation of protein and proportion of clustered beads were significantly lower (Figure 4H-J), supporting our conclusion that these mutations interfere with DNA binding. Thus, efficient DNA-binding of ZYG-9 promotes its superficial accumulation around DNA beads and its ability to cluster them.

### ZYG-9’s arginine patch is required for tight clustering of chromosomes and packaging them into polar bodies

To test if ZYG-9-DNA binding has a pivotal role in egg formation, we engineered homozygous oocytes expressing GFP::ZYG-9^5RG^ from the endogenous genomic locus (Figure 4A). During metaphase I, GFP::ZYG-9^5RG^ efficiently accumulated at correctly formed bipolar spindles (Figure 5A-5C). Sister chromosomes pushed apart from each other normally during anaphase I (Figure 5B). These results indicate that DNA binding of ZYG-9 is not required for its spindle-organizing activity throughout meiosis I. However, 5RG oocytes experienced chromosome dispersal during anaphase and beyond, indicated by a significant increase in the lateral inter-chromosomal distance compared with WT oocytes (Figure 5D). Moreover, ∼10% of 5RG oocytes failed to package the correct number of chromosomes into the first polar body (Figure 5D and 5I); in these cases, chromosomes underwent regression from the polar body back to the egg cytoplasm 5 min after chromosome segregation (yellow arrow heads, Figure 5D; Video S3 and S4). These chromosome abnormalities were more frequent during anaphase II (28.6% occurrence, Figure 5E-5G). Thus, ZYG-9 binding to DNA is required to keep chromosomes tightly clustered and confined into the limited space of the polar body during meiotic completion.

**Figure 5.**
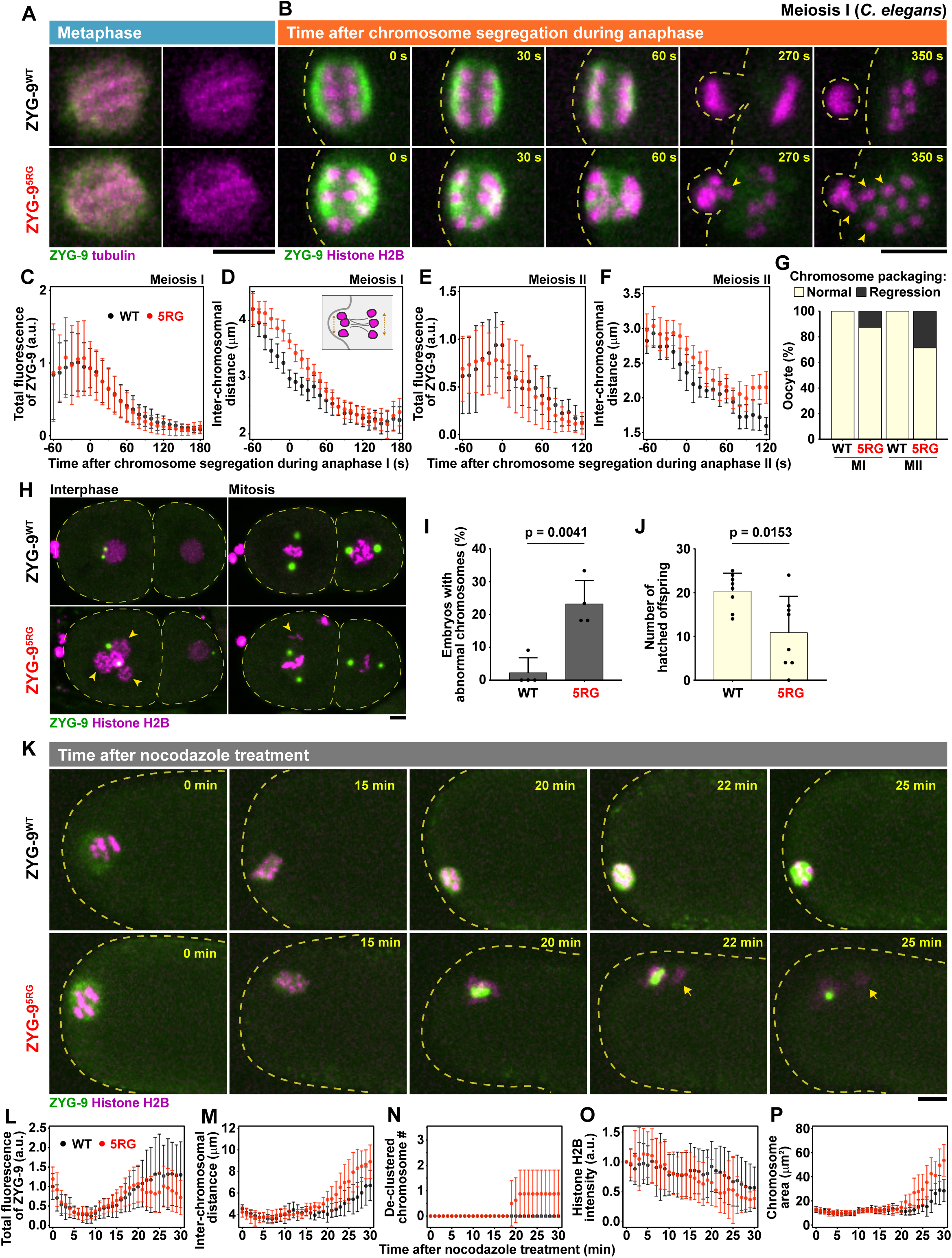
The ZYG-9 arginine patch ensures tight chromosome clustering to prevent extra chromosome carryover into eggs. **(A)** Co-visualization of GFP::ZYG-9 (green) and mCherry::tubulin (magenta) during metaphase I (left). Morphology of the spindle indicated by the tubulin signal (right). **(B)** Time-lapse imaging of GFP::ZYG-9 (green) and mCherry::histone H2B (magenta) throughout anaphase I. Yellow allow heads indicate regressed chromosomes from the polar body to the oocyte cytoplasm. **(C-G)** Normalized total fluorescence of spindle-associated GFP::ZYG-9 (C, E), lateral distance of the farthest chromosomes (D, F), and population of oocytes with regressed chromosomes (G) during anaphase I and II. Data are mean ± 95% C.I. (WT, n= 20; 5RG, n= 23). **(H)** Co-visualization of GFP::ZYG-9 (green) and mCherry::histone H2B (magenta) during interphase (left) and mitosis (right) in 2-cell stage embryos. Yellow arrowheads indicate extra nucleus and ectopic chromosomes. **(I)** Population of 1-4 cell stage embryos with abnormal chromosomes in (H). Data represent mean ± SD from four independent experiments (WT, n= 63; 5RG, n= 72). p-value was determined by unpaired t-test. **(J)** Number of hatched eggs from single mother expressing GFP::ZYG-9^WT^ (n= 8) or GFP::ZYG-9^5RG^ (n= 8). Data represent mean + SD. p-value was determined by unpaired t-test. **(K)** Time-lapse imaging of *ex utero* oocytes treated with 20 µM nocodazole. Yellow arrows indicate de-clustered chromosomes. **(L-P)** Quantification of total fluorescence of GFP::ZYG-9 condensate (L), distance between the farthest chromosomes (M), number of de-clustered chromosomes (N), normalized mCherry::Histone H2B intensity (O), and chromosome size (P) after nocodazole treatment. Data are mean ± 95% C.I. (WT, n= 8; 5RG, n= 10). All scale bars are 5 µm.

We next tested if improper retention of chromosomes in the oocytes affected embryo development. Unlike WT embryos, 5RG embryos displayed chromosomal abnormalities. 5RG embryos frequently formed multi-nucleated cells during interphase (Figure 5H). Mitotic cells in 5RG embryos also showed ectopically localized chromosomes, where chromosomes are outside the spindle region (Figure 5H). Thus, DNA binding activity of ZYG-9 is required to prevent aneuploidy in embryos. The frequency of 5RG embryos with these abnormal karyotypes (23.4%, Figure 5J) was comparable with occurrence of the chromosome regression during meiosis II (28.6%, Figure 5G), suggesting that extra chromosomes are carried over from the oocyte into the mitotic embryo. Consistent with this observation, the 5RG mutation significantly reduced overall fertility, measured by the number of successfully hatched offspring (Figure 5A and 5B). Thus, ZYG-9’s function in packaging chromosomes during meiosis helps ensure the formation of viable euploid eggs.

We previously showed that ZYG-9 is required for chromosome clustering in the absence of the spindle (Figure 2F). To test if this activity depends on ZYG-9’s DNA-binding ability, we treated WT and 5RG oocytes with nocodazole during metaphase I. WT oocytes formed perichromosomal ZYG-9 condensates that stably tethered chromosomes in the spindle-less oocytes, as expected (Figure 5K). In contrast, GFP::ZYG-9^5RG^ formed unstable condensates; fluorescent signal of the superficial ZYG-9 declined 20 min after nocodazole treatment (Figure 5K and 5L), suggestive of defective condensation, consistent with our *in vitro* experiments (Figure 4F). Chromosomes subsequently drifted apart and de-compacted in the 5RG oocytes (Figure 5M-5P), as seen after acute depletion of ZYG-9 in anaphase (Figure 2A). Our results demonstrate that the arginine-rich motif of ZYG-9 is required for its condensation around chromosomes, which keeps them stably corralled and compacted both in the presence or absence of the meiotic spindle.

## Discussion

In summary, we identified that the protein ZYG-9 surrounds meiotic chromosomes and spatially constrains them throughout anaphase in *C. elegans* oocytes. This activity is likely independent of ZYG-9’s canonical role as a microtubule polymerase (35, 37, 41, 42) for two reasons: 1) ZYG-9 encapsulates and traps chromosomes even after the meiotic spindle is disassembled with nocodazole, and 2) five point mutations in an IDR (5RG mutant)—outside of any characterized microtubule or tubulin-binding region—cause chromosome dispersal without affecting meiotic spindle morphology or function. Our results suggest that liquid-like condensates of ZYG-9 serve as chromosome glue when the meiotic spindle undergoes architectural rearrangement and pushes chromosomes toward the cell periphery for expulsion into polar bodies (Figure 6). ZYG-9 could also corral meiotic chromosomes in times of spindle stress which could be triggered, in theory, by exposure to extreme cold or environmental poisons, both of which are known to depolymerize microtubules (43–45). It is possible that this organizing principle is conserved since ZYG-9 belongs to a highly conserved family of proteins that contain tandem TOG domains. In support of this idea, we found that chTOG localizes around meiotic chromosomes in mouse oocytes, even in the absence of the spindle (Figure S4). Furthermore, overexpressed domains of Msps (fly homolog) and full-length XMAP215 (frog homolog) can accumulate at chromosomes (46, 47). Future work is required to clarify the functional roles of TOG family proteins in genome organization in these species.

**Figure 6.**
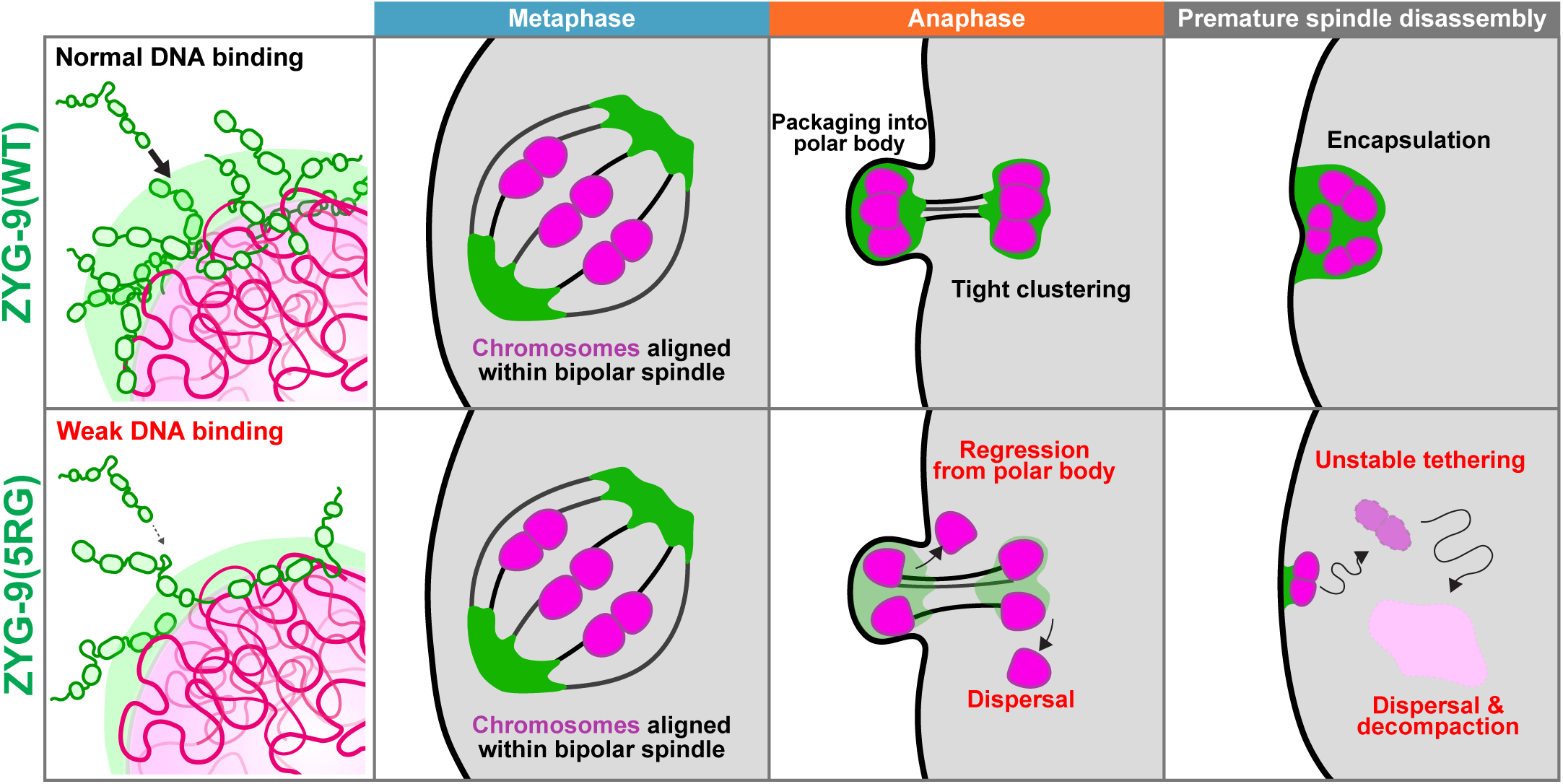
Model: Surface-acting glue spatially confines chromosomes during female meiosis. ZYG-9/chTOG (green) directly binds DNA (magenta) via an arginine-rich IDR. Top Panels: During anaphase, wild-type ZYG-9 surrounds meiotic chromosomes, acting as a superficial glue to enable tight chromosome clustering and packaging into the small polar body. In the absence of the spindle, ZYG-9 condenses into a thick phase that encapsulates all meiotic chromosomes to keep them clustered and compacted. Bottom Panels: Decreasing DNA affinity of ZYG-9 (5RG mutant) does not affect bipolar spindle formation and metaphase chromosome alignment. However, in 5RG oocytes, anaphase chromosomes undergo dispersal and regression from the polar body to the oocyte cytoplasm, leading to aneuploidization and infertility. In the spindle-less 5RG oocyte, ZYG-9 coating of chromosomes is destabilized, and chromosomes de-cluster and de-compact.

Non-cytoskeletal, surface-acting agents have been reported to assist the organization of mitotic chromosomes in somatic cells (48). Human Ki-67 coats mitotic chromosomes, acting first as a physical barrier to prevent chromosome tangling in mitosis, then as a glue to keep chromosomes close during mitotic exit (49–53). However, a function for Ki-67 has not been reported in meiosis, nor does this protein exist outside of vertebrates (54–56). Barrier to Autointegration Factor (BAF) tethers chromosomes to each other and to the nuclear envelope (57, 58), yet its depletion in *C. elegans* oocytes had no effect on meiotic chromosome organization (59, 60). Thus, prior to our work, it was unclear if surface-acting agents are used to organize meiotic chromosomes, which differ in architecture from mitotic chromosomes. Our data suggest that a dynamic, liquid-like spindle pole is essential for coating chromosomes during a vulnerable time to prevent their dispersal in a vast oocyte cytoplasm (Figure 6). Thus, binding of chromosomes with a surface-acting glue may be a general organizing principle for the genome across species and cell types. Furthermore, our findings could be important for understanding the molecular basis of meiotic errors and developmental syndromes in the eggs from aging or infertile women.

## Supporting information

Video 1. Localization of spindle pole proteins (green) after nocodazole treatment.

Video 2. Reconstituted chromatin droplets (magenta) with ZYG-9 (green)

Video 3. Meiosis in GFP::ZYG-9(WT) oocyte.

Video 4. Meiosis in GFP::ZYG-9(5RG) oocyte.

## Acknowledgements and Author Contributions

We thank Sarah Wignall, Bruce Bowerman, and Jessica Feldman for providing strains; the Quantitative Light Microscopy Core Facility for help with super-resolution microscopy; and the Macromolecular Biophysics core for help with mass photometry. K. Yaguchi performed and analyzed all experiments; L. Chen prepared reconstituted chromatin; N. Familiari expressed proteins; J. Tafur helped with strain construction and worm maintenance; A. Mohammadi-Sangcheshmeh and E. Grow prepared mouse oocytes; M. Rosen and J. Woodruff supervised the project; K. Yaguchi and J. Woodruff wrote the paper with input from all authors.

## Funding

K. Yaguchi was supported by a Human Frontier Science Program Fellowship (LT0064/2022-L). J.B. Woodruff was supported by a Welch Foundation Grant (V-I-0004-20230731), and R35 grant from the National Institute of General Medical Sciences (1R35GM142522), and the Endowed Scholars program at UT Southwestern. Research in the Rosen lab was supported by the Howard Hughes Medical Institute and the NIH (R35GM141736). EJG is supported by NIH (R35GM159821) and CPRIT (RR210077) grants. Some data presented in this report were acquired with a mass photometer that was supported by award S10OD030312-01 from the National Institutes of Health.

## MATERIALS AND METHODS

### Experimental model and subject details

*C. elegans* worm strains were grown on nematode growth medium (NGM) plate at 20°C, following standard protocols (http://www.wormbook.org). Worm strains used in this study are listed in Table S1. Plasmids used for RNAi feeding and protein expression are listed in Tables S2. Oligo nucleotides used for CRISPR strain generation and *in vitro* DNA assay are listed in Table S3. For expression of recombinant proteins, we used suspended SF9-ESF *Spodoptera frugiperda* insect cells grown at 27°C in ESF 921 Insect Cell Culture Medium, Protein-Free (Expression Systems), supplemented with FBS (2% final concentration; (61)) or *Escherichia coli* bacteria cells (BL21(DE3) CodonPlus-RIL) grown in LB medium containing 100 µg/ml kanamycin. For obtaining mouse oocytes, BDF1 mice were maintained in the UTSW Cecil H. and Ida Green Center.

### Live imaging of *C. elegans* oocytes

For *in utero* imaging, adult worms were placed in a droplet of 5 µl meiosis medium (62) containing 0.02% levamisole and mounted with 5% agar pad on a cover glass. For *ex utero* imaging, adult worms were dissected in 5 µl meiosis medium on a cover glass. Before mounting on the microscope stage, the meiosis medium droplet was covered by mineral oil (SIGMA) to prevent evaporation. Time-lapse images were taken using an inverted Nikon ECLIPSE Ti2 equipped with Yokogawa Spinning Disk Field Scanning Confocal System (CSU-W1), piezo Z stage, and iXon Ultra EMCDD camera (Andor) controlled by NIS-Elements software. On this system, simultaneous dual-color imaging with 488- and 561-nm laser was achieved using OptoSplit II beam splitter (Cairn). For most experiments, we used a 100× 1.35-NA Plan Apochromat silicone oil objective to acquire 5-10 × 1-2 µm Z-stacks. Super-resolution images were taken using an inverted Nikon ECLIPSE Ti2 equipped with Yokogawa Spinning Disk system (Super Resolution by Optical Pixel Reassignment System; CSU-W1 SoRa), and ORCA-Fusion CMOS camera (HAMAMATSU) controlled by NIS-Elements software. To investigate the spatial association between chromosomes and spindle poles, mCherry::H2B and GFP::ZYG-9 were visualized using a 60× 1.42-NA Plan Apochromat oil objective with 4× intermediate magnification.

### Time-controlled perturbation of the spindle pole and the spindle microtubules

For acute degradation of degron::GFP::ZYG-9 during meiotic progression, 2× auxin-containing 5 µL meiosis medium was injected into the culture on the microscope stage (10 mM final concentration). As a vehicle control, 0.5% of ethanol in meiosis medium was used. For depolymerization of the spindle microtubules during metaphase, *ex utero* oocytes were treated with 20 µM nocodazole in meiosis medium. mCherry::TBB-2 was co-visualized with the spindle pole proteins to confirm that the spindle was disassembled by nocodazole treatment (Video S1). For degradation of degron::GFP::ZYG-9 in nocodazole-arrested *ex utero* oocytes, oocytes were first treated with 20 µM nocodazole in 5 µL meiosis medium for 15 min, then 2× auxin-containing 5 µL meiosis medium was injected into the medium (500 µM final concentration) as described in Figure 3A.

### RNAi

*Ndc-80 RNAi* was done by feeding using a feeding clone (Table S2). Briefly, bacteria expressing *ndc-80* dsRNA were grown at 37°C in LB medium containing 100 µg/mL ampicillin until the culture reached an OD600 at 0.6, followed by the induction with 1 mM IPTG supplementation for 2 hr to ensure the dsRNA expression. The pre-induced culture was seeded onto NGM plates supplemented with 1 mM IPTG and 100 µg/mL ampicillin. Plates were incubated at 37°C for 4 hr. To deplete NDC-80 protein in oocytes, L4 hermaphrodites were grown on RNAi plates at 20°C for 24-30 hr before imaging.

### Protein purification

Full length proteins were expressed in *Spodoptera frugiperda* insect cells using the FlexiBAC baculovirus plasmids (Table S2)(61). Cells were lysed by dounce homogenization in KCl buffer (25 mM Hepes, pH 7.4, 300 mM KCl, 20 mM imidazole, 1 mM PMSF, 0.1% CHAPS) or NaCl buffer (50 mM Tris-HCl, pH 8.0, 200 mM NaCl, 3% Glycerol, 1 mM PMSF, 0.1% CHAPS), and extracts were clarified by centrifugation for 30 min at 28000 rpm (BECKMAN 50.2Ti). To purify GFP::ZYG-9 and unlabeled TAC-1, the clarified lysates with a KCl buffer were loaded to columns packed with Ni-NTA resin (QIAGEN), and the protein-bound beads were washed with a lysis buffer supplemented with 200 mM KCl (500 mM final concentration). Proteins were eluted with 150 mM imidazole. SPD-5 was purified using MBP-trap beads (ChromoTek), as previously described (63). Porcine tubulin was purified and stored as previously described (64). ZYG-9 fragments were expressed in *E. coli* (BL21(DE3) CodonPlus-RIL) induced with 0.3 mM IPTG at 16°C for overnight or with 0.8 mM IPTG at 28°C for 4 hr. Cell pellets were frozen in liquid nitrogen in a KCl buffer. Cells were lysed using an Emusiflex followed by sonication. Proteins were then purified following the same procedure as for GFP::ZYG-9. To purify unlabeled ZYG-9 and mScarlet::ZYG-9, the clarified lysates with a NaCl buffer were loaded onto columns packed with Strep-TactinXT resin (iba). Protein was eluted through on-column cleavage with with 10 µg/mL 3C protease. To remove 3C protease, eluates were passed over Ni-NTA resin. All eluates were concentrated with 30 or 100 K Amicon Ultra Centrifugal Filter (Millipore). Concentration of purified proteins was measured using a Nanodrop (ThermoFisher Science) and stored in a storage buffer (25 mM Hepes, pH 7.4, 500 mM KCl, and 10% glycerol) at -80°C.

### DNA binding assay

50-bp dsDNA substrate was prepared by annealing complementary oligonucleotides (poly-A and poly-T sequences; IDT). Poly-A oligonucleotide was labeled with 6-FAM at the 5’ end. Successful annealing was confirmed by electrophoresis. Linear 5-kbp dsDNA was prepared by PCR amplification of a plasmid sequence (Table S3), followed by gel-purification. EMSA reaction was assembled with a final concentration of 20 pM 5-kbp dsDNA or 200 nM 50-bp dsDNA and varying concentration of ZYG-9 proteins in 25 mM Hepes pH 7.4, 200 mM KCl, 2 mM MgCl_2_, 2 mM DTT; followed by incubation for 3 hr at RT. 10% glycerol was added before loading into EMSA gels. DNA-protein complexes were resolved by electrophoresis at RT on 2% TBE-agarose gel (60 min at 10 V/cm) for the 50-bp dsDNA substrate, and at 4°C on 0.7% 0.5× TAE-agarose gel (200 min at 10 V/cm) for the 5-kbp dsDNA substrate, respectively. 5-kbp dsDNA was detected by SYBR Gold staining (Invitrogen). Apparent equilibrium dissociation constant K_d_^app^ was defined with the fitted binding curves in Figure 4.

### Mass photometry

200 nM ZYG-9 protein in PBS (SIGMA) was centrifuged and supernatant was collected to remove potential aggregates before the analysis. Protein was ten-times diluted on a cleaned glass coverslip (20 nM final concentration), and analyzed by a TwoMP mass photometer (Refeyn, UTSW Macromolecular Biophysics Resource). Molecular mass was calibrated and analyzed using a standard curve of four species of BSA (monomer, dimer, trimer, tetramer).

### *in vitro* condensate assays

ZYG-9 self-assemblies and co-condensation with DNA were assembled by diluting 5× protein/DNA solution in a physiological salt buffer (25 mM Hepes, pH 7.4, and 100 mM KCl). The mixtures were immediately placed in PEGylated glass-bottom 96-well dishes (Corning, 4850, high content imaging dish) and incubated for 30 min at RT. The imaging plates were prepared by incubation with 5% Hellmanex (HellmaAnalytics) at 37°C for 4 hr and etching with 1 M NaOH at RT for 30 min, followed by incubation with 20 mg/mL 5K mPEG-silane (PEGWorks) in 95% ethanol at RT for 18 hr. PEGylated glasses were washed with 95% ethanol and deionized-distilled water, as previously described (32). Proteins assemblies were imaged on an inverted Nikon ECLIPSE Ti2 equipped with an AX modular confocal system controlled by NIS-Elements software. Co-existing chromatin droplets were formed by co-incubation of 500 nM GFP::ZYG-9 and 500 nM chromatin at RT for 30 min in a nucleosome buffer (25 mM Tris-OAc, pH 7.5, 80 or 150 mM KOAc, 1 mM MgOAc, 0.1 mM EDTA, 5 mM DTT, 0.1 mg/mL BSA, 5% glycerol, 2 µg/mL glucose oxidase, 350 ng/mL catalase). The co-existing condensates were imaged using a Leica SP8 confocal microscope equipped with a resonant scanning stage and a hybrid (HyD) detector controlled by Leica X software.

### DNA beads assay

Alexa 647-labeled 1-kbp DNA was prepared by PCR amplification of a plasmid sequence using primers labeled with Alexa647 (forward) and biotin (reverse) at 5’ end of each (IDT), followed by gel-purification (Table S3). DNA beads were prepared by incubating 1.5 mg of the gel-purified DNA with 1 mg of 2.8 µm diameter Streptavidin beads (M-270, Invitrogen) at 4°C overnight in 50 µl of a binding buffer (10 mM Hepes pH 7.4, 0.5 mM EDTA, 1 M NaCl). DNA-bound beads were washed with a binding buffer supplemented with 2 M NaCl and stored in 25 mM Hepes, pH 7.4. DNA beads were incubated in a physiological salt buffer for 10 min at RT while shaking at 300 rpm in the absence or presence of 500 nM mScarlet::ZYG-9 proteins in the reaction. The reaction mixture was gently moved into the imaging plate for the image aquisition and analysis.

### Immunostaining of mouse oocytes

BDF1 mice were injected with 5 IU pregnant mare serum gonadotropin, and GV oocytes were obtained into M2 medium ∼48 hr later. Cumulus cells were removed in the culture by manual pipetting. For immunostaining of MI oocytes, oocytes were fixed with 4% PFA in DPBS for 20 min at 37°C at 8 hr after *in vitro* maturation. For immunostaining of nocodazole-treated oocytes, oocytes were treated with 20 µM nocodazole for 15 min at 5 hr after *in vitro* culture, followed by 4% PFA fixation. Fixed oocytes were permeabilized with 0.5% triton X-100 in DPBS at 4°C overnight, followed by treatment with BSA blocking buffer (150 mM NaCl, 10 mM Tris-HCl, pH 7.5, 5% BSA, and 0.1% Tween 20) for 1 hr at 25°C. Samples were incubated with a primary antibody (rabbit poly-clonal anti-chTOG, ab236981, 1:200) at 4°C overnight, and then incubated with secondary antibodies (Alexa647-conjugated Donkey anti-Rabbit, A31572, 1:200; FITC-conjugated mouse monoclonal anti-a-tubulin, B-5-1-2, 1:1000) at 4°C overnight. After each treatment, oocytes were washed three times with DPBS. Stained oocytes were mounted with a mounting medium containing propidium iodide (VECTASHIELD) and imaged on an inverted Nikon ECLIPSE Ti2 equipped with Yokogawa Spinning Disk Field Scanning Confocal System (CSU-W1).

### Image quantification and statistical analysis

The fluorescent intensities were measured using a line tool manually or a semi-automated segmentation with the subtraction of background signals in Fiji (NIH). The organelle-organelle distances were measured between the points of their highest intensities. Inter-chromosomal distance was measured between the farthest chromosomes in oocytes in Figure 2 and 5. Total fluorescence of protein signal was defined as the mean intensity multiplied by its area. This value was displayed in arbitrary units (a.u.). For all time-lapse imaging data, mean values and 95% C.I.s were plotted using GraphPad Prism. Analyses for significant difference between two groups were conducted using unpaired *t* test in Figure 4 and 5.

## SUPPLEMENTAL MATERIAL

### Supplemental Figure Legends

**Figure S1.**
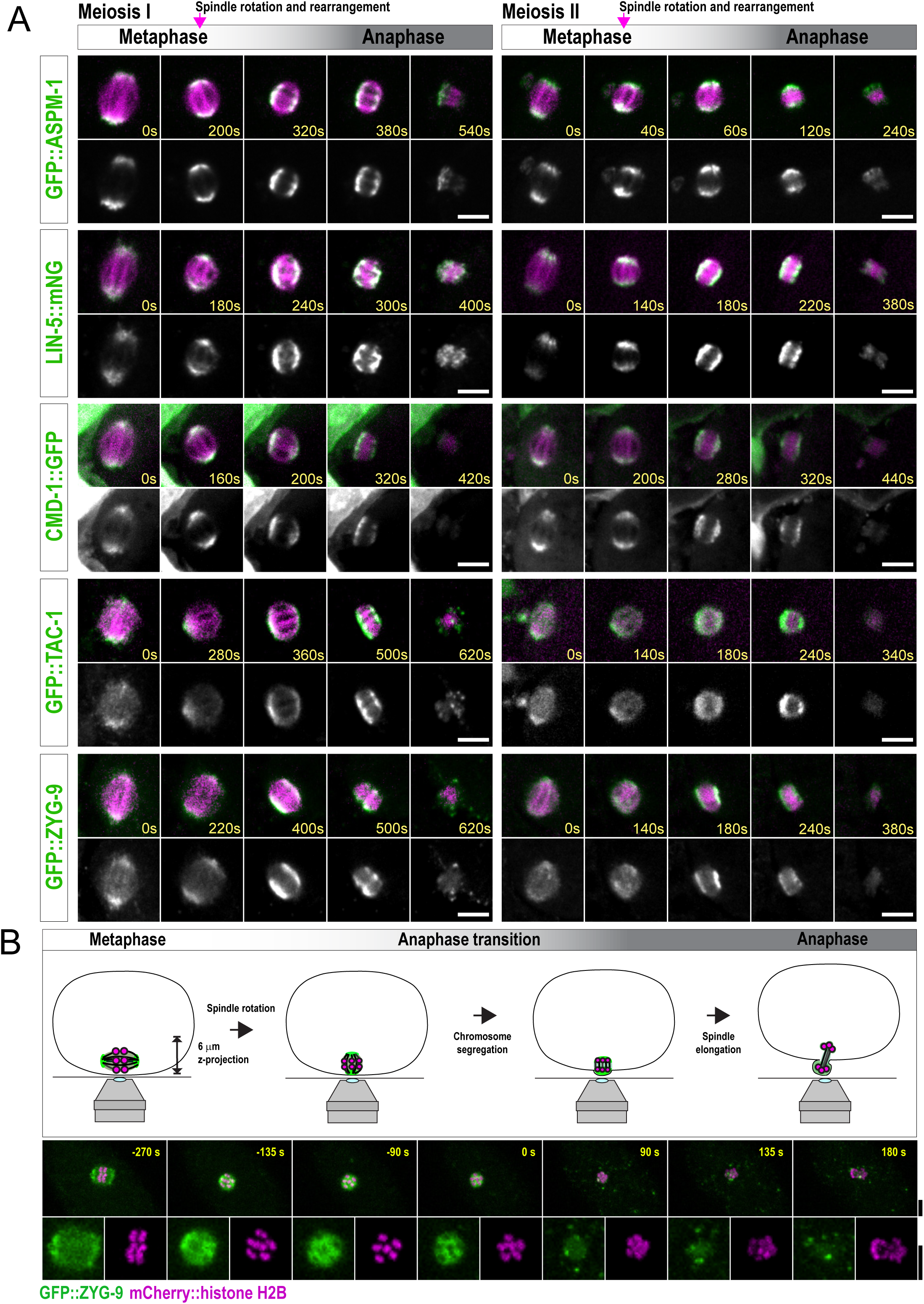
Translocation of spindle pole proteins during the anaphase transition in *C. elegans* meiosis. **(A)** Time-lapse imaging of *in utero* oocytes during meiosis I (left) and II (right). Each spindle pole protein was endogenously labeled with GFP or mNeonGreen (green). mCherry::tubulin was co-visualized to monitor spindle shortening and rotation during the anaphase transition (magenta). Spindle shortening was set as 0 s. **(B)** Diagram of *ex utero* oocytes forming the first polar body facing toward the microscope objective (top). GFP::ZYG-9 (green) encapsulated chromosomes indicated by mCherry::H2B (magenta) during the anaphase transition (middle). Enlarged images of each signal show voids of GFP::ZYG-9 signals occupied by chromosomes at the anaphase transition (bottom). Onset of chromosome segregation was set as 0 s. All scale bars are 5 µm.

**Figure S2.**
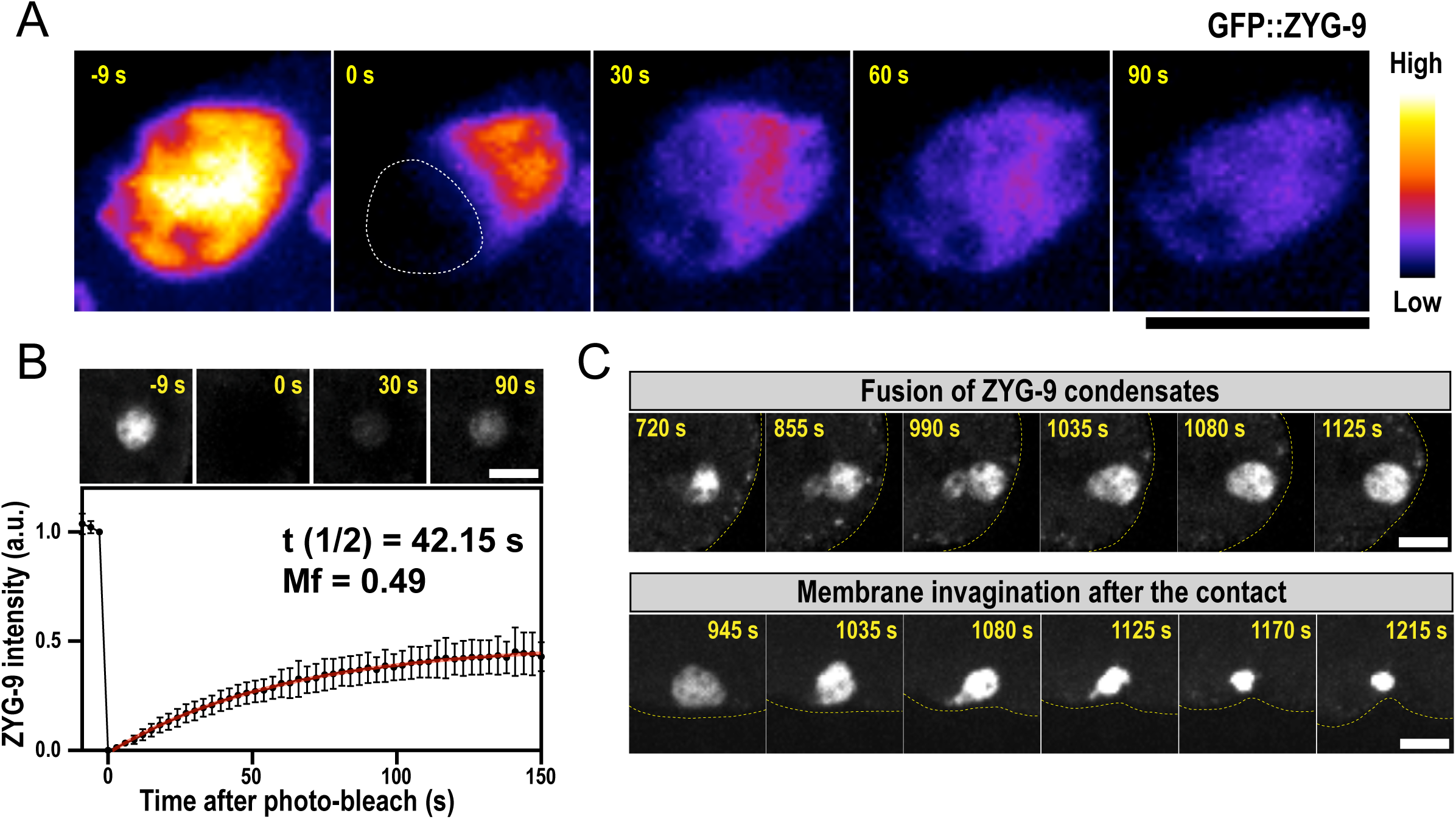
ZYG-9 forms into a micron-scale, liquid-like condensate in the nocodazole-treated oocyte. **(A)** Fluorescence recovery after partial photobleaching of GFP::ZYG-9 condensates in nocodazole-treated *ex utero* oocytes. The photobleached region recovered fluorescent signal coincident with a decrease in the intensity of the unbleached region. Images are pseudo-colored to improve the contrast (high intensity to low intensity: white, yellow, red, purple, blue, black). **(B)** Fluorescence recovery after full photobleaching of GFP::ZYG-9 condensates in nocodazole-treated oocytes. Data is mean ± 95% C.I. (n= 10). Data were fitted with a single exponential function to obtain the mobile fraction (Mf, red line). **(C)** Coalescence and wetting behavior of GFP::ZYG-9 in nocodazole-treated oocytes. Broken lines indicate the membrane. All scale bars are 5 µm.

**Figure S3.**
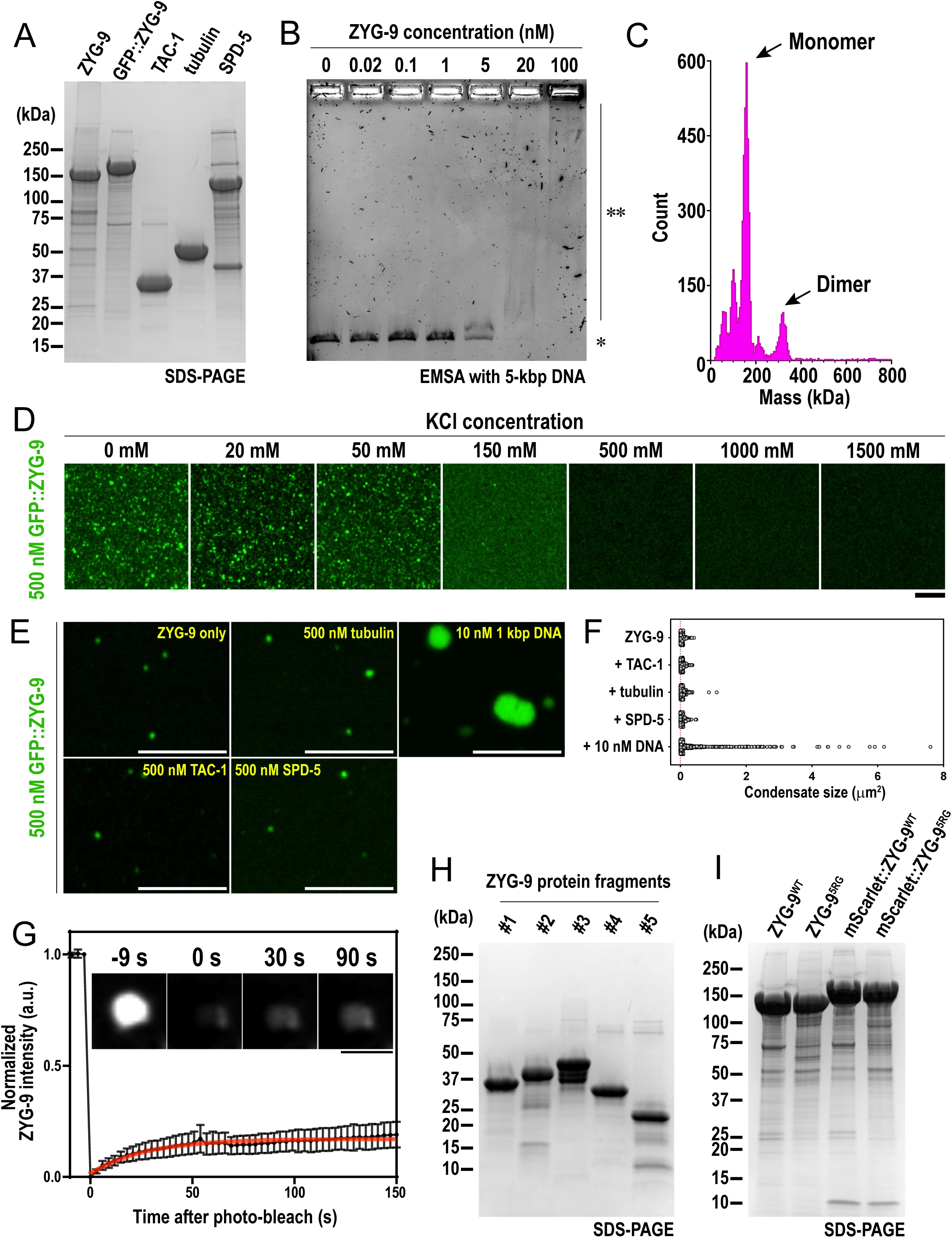
ZYG-9 and DNA co-condense *in vitro*. **(A)** SDS-PAGE of purified proteins. 10 µM of each protein was loaded into the gel with 5 µL/well. **(B)** Electrophoretic mobility shift assay of 5 kbp dsDNA co-incubated in the presence of 0-100 nM ZYG-9. Unbound (*) and bound (**) species are indicated. **(C)** Mass photometric analysis of 20 nM ZYG-9 proteins in PBS solution. Monomeric and dimeric peaks are indicated. **(D)** Confocal images of 500 nM GFP::ZYG-9 incubated under different salt concentrations. **(E)** Confocal images of GFP::ZYG-9 co-incubated with different molecules. **(F)** Size measurement of GFP::ZYG-9 mass in (E). Each data point represents area of single punctum/condensate from two independent experiments. **(G)** Fluorescence recovery of GFP::ZYG-9 after photobleaching the co-condensates containing 1 kbp DNA *in vitro*. The data were fitted with a single exponential function. All scale bars are 5 µm. **(H and I)** SDS-PAGE of ZYG-9 protein fragments purified from bacteria cells (H) and full length ZYG-9 recombinant proteins purified from insect cells (I). 10 µM of each protein was loaded into the gel with 10 µL/well.

**Figure S4.**
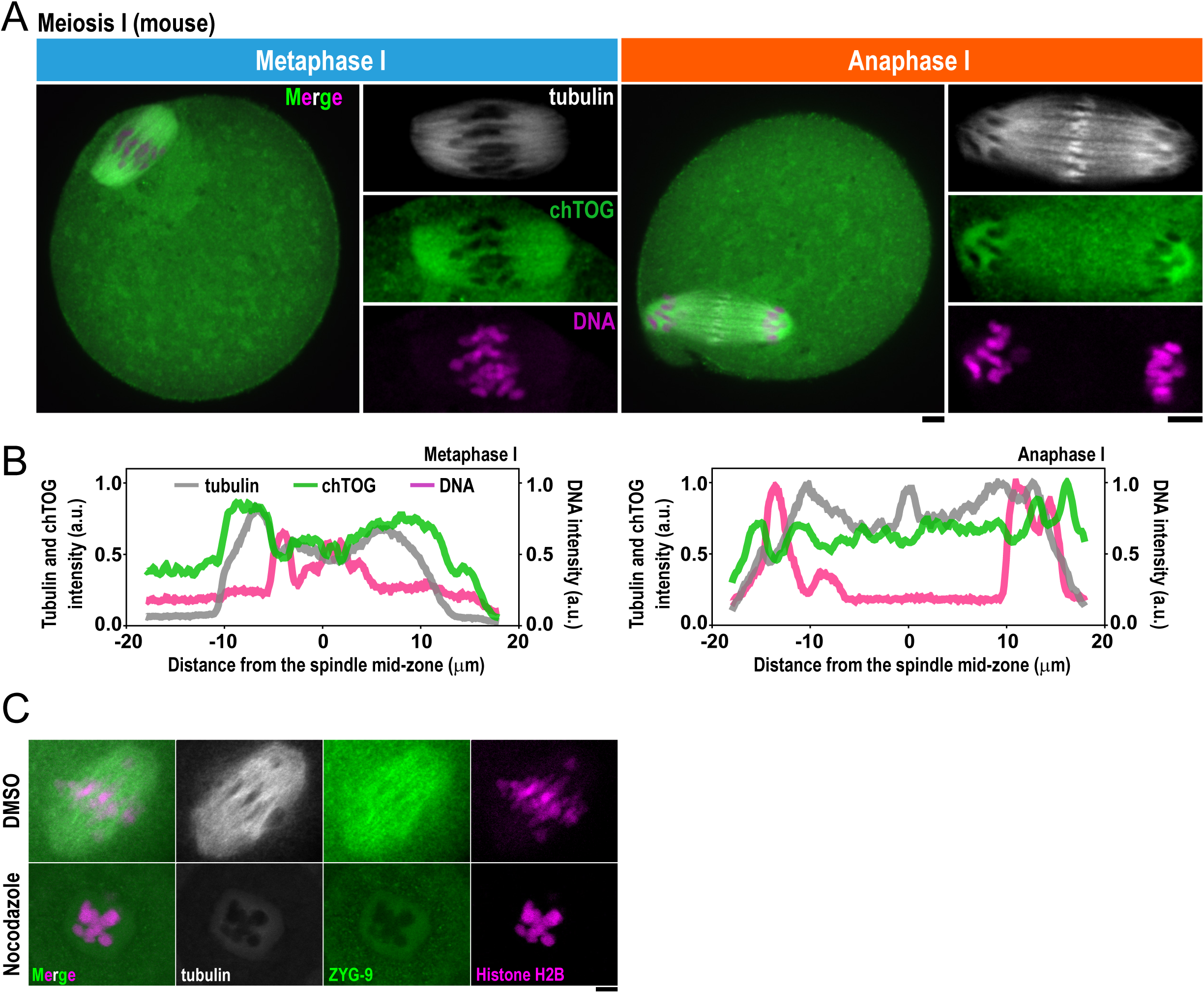
chTOG associates with chromosomes in mouse oocytes. **(A)** Immunostaining of tubulin (white), chTOG (green), and DNA (magenta) during metaphase I and anaphase I in mouse oocytes (n= 14). **(B)** Line scans of signal intensity across the long axis of the spindle. Data are normalized to maxima of each signal. **(C)** Mouse MI oocytes treated with DMSO (n= 13) or 20 µM nocodazole (n= 13) for 15 min. All scale bars are 5 µm.

**Table S1:** *C. elegans* strains used in this study.

| Strain # | Genotype |
| --- | --- |
| N2 | Wild-type (ancestral) |
| EU3169 | zyg-9(or1956[gfp::zyg-9]) II. |
| PHX10780/<br>JWW 270 | zyg-9(or1956[gfp::zyg-9(R895G, R896G, R900G, R902G, R904G)]) II (this study). |
| JWW 278 | zyg-9(or1956[gfp::zyg-9]) II; ltlIs37 [(pAA64) pie-1p::mCherry::his-58 + unc-119(+)] IV. |
| JWW 279 | zyg-9(or1956[gfp::zyg-9(R895G, R896G, R900G, R902G, R904G)]) II; ltlIs37 [(pAA64) pie-1p::mCherry::his-58 + unc-119(+)] IV. |
| SMW24 | Pzyg-9::AID::EmGFP::zyg-9 (C6->T—PAM site mutation) II; unc-119(ed3); ieSi38 [Psun-1::TIR1::mRuby::sun-1 3'UTR, cb-unc-119(+)] IV |
| JLF105/WOW12 | zyg-9[zf::GFP::ZYG-9] II |
| JWW198 | zyg-9[zf::GFP::ZYG-9] II; [pJA138; 5'pie-1::mCherry::tbb-2::3'pie-1] (JA1559) |
| JWW257 | Pzyg-9::AID*::EmGFP::zyg-9 (C6->T—PAM site mutation) II; ltlIs37 [(pAA64) pie-1p::mCherry::his-58 + unc-119(+)] IV; wrdSi50[mex-5p::TIR1::F2A::mTagBFP2::AID*::NLS::tbb-2 3'UTR] (I:-5.32). |
| JWW183 | vie11[pAD676; gfp::tac-1]II (DAM858); [pJA138; 5'pie-1::mCherry::tbb-2::3'pie-1] (JA1559) |
| JWW208 | cmd-1[cmd-1::eGFP] V (this study); [pJA138; 5'pie-1::mCherry::tbb-2::3'pie-1] (JA1559) |
| JWW194 | lin-5(cp288[lin-5::mNG-C1^3xFlag]) II (LP585); [pJA138; 5'pie-1::mCherry::tbb-2::3'pie-1] (JA1559) |
| JWW182 | aspm-1(or1935[GFP::aspm-1]) I (EU2861); [pJA138; 5'pie-1::mCherry::tbb-2::3'pie-1] (JA1559) |

**Table S2:** Plasmids used in this study.

| Vector type | Gene/Insert | Identifiers |
| --- | --- | --- |
| Protein expression (pOCC) | eGFP:: ZYG-9 (wt)::6xHis | pKY16 |
| Protein expression (pOCC) | MBP::PreScission::SPD-5::PreScission::6xHis | TH0861 |
| Protein expression (pOCC) | 6xHis::TAC-1 | TH1034 |
| Protein expression (pET) | ZYG-9 (Fragment #1, a.a. 1-280)::6xHis | pKY24 |
| Protein expression (pET) | ZYG-9 (Fragment #2, a.a. 281-598)::6xHis | pKY25 |
| Protein expression (pET) | ZYG-9 (Fragment #3, a.a. 599-975)::6xHis | pKY26 |
| Protein expression (pET) | ZYG-9 (Fragment #4, a.a. 976-1265)::6xHis | pKY27 |
| Protein expression (pET) | ZYG-9 (Fragment #5, a.a. 1241-1415)::6xHis | pKY28 |
| Protein expression (pOCC) | StrepII::3C PreScission::ZYG-9 (wt) | pKY47 |
| Protein expression (pOCC) | StrepII::3C PreScission::ZYG-9 (5RG) | pKY73 |
| Protein expression (pOCC) | StrepII::3C PreScission::mScarlet::ZYG-9 (wt) | pKY39 |
| Protein expression (pOCC) | StrepII::3C PreScission::mScarlet::ZYG-9 (5RG) | pKY74 |
| RNAi feeding clone | *ndc80* fragment (exon 2-6) | JWB215 |
| CRISPR donor | eGFP | pJJR82 |

**Table S3:**
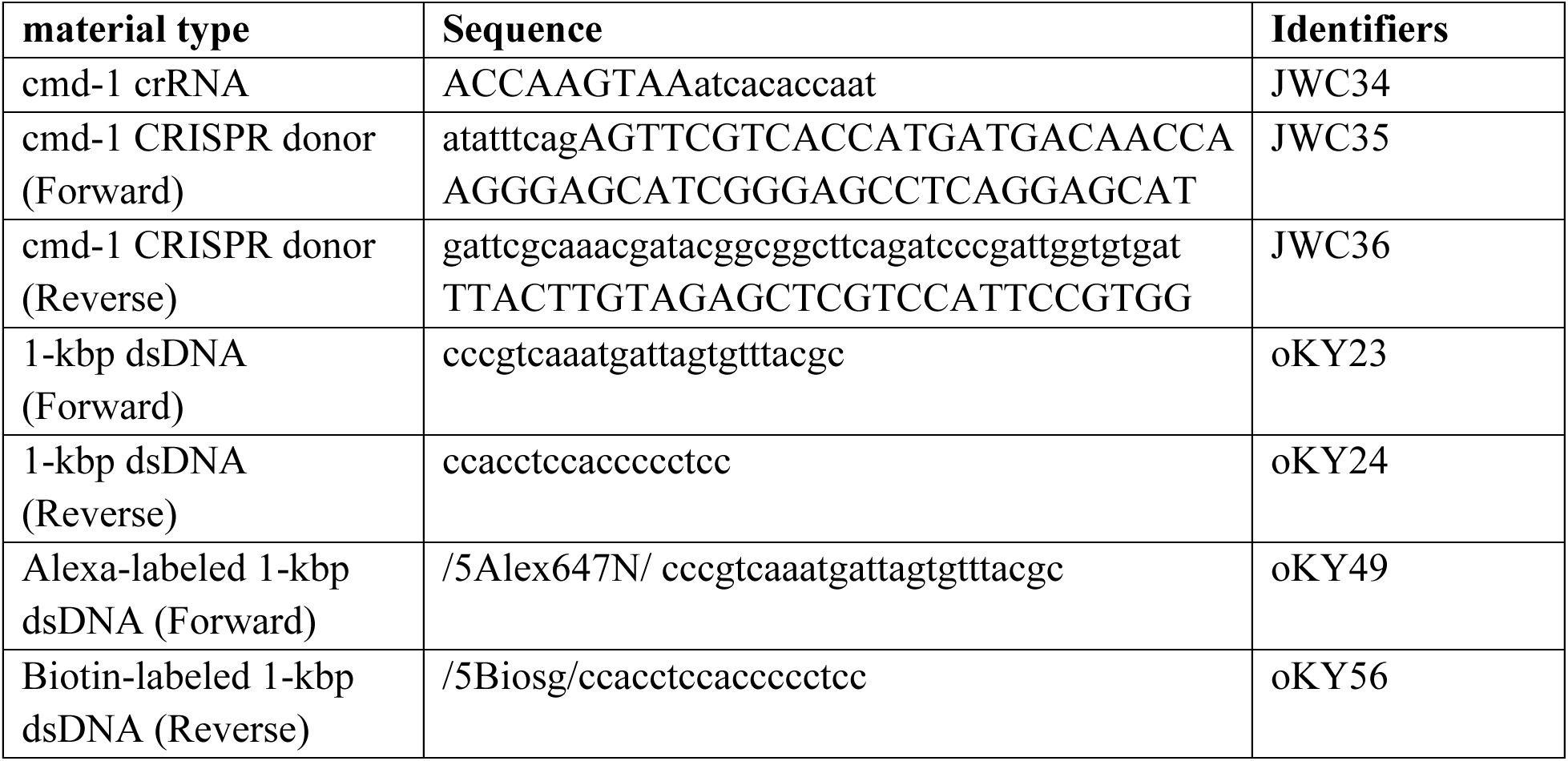
Oligonucleotides used in this study.

## Notes

### Competing Interest Statement

The authors have declared no competing interest.

## REFERENCES

1. Kyogoku H, Kitajima TS. The large cytoplasmic volume of oocyte. J Reprod Dev. 2023;69(1):1–9.

2. Goyanes VJ, Ron-Corzo A, Costas E, Maneiro E. Morphometric categorization of the human oocyte and early conceptus. Hum Reprod. 1990;5(5):613–8.

3. Klatsky PC, Carson SA, Wessel GM. Detection of oocyte mRNA in starfish polar bodies. Mol Reprod Dev. 2010;77(5):386.

4. Evans JP, Robinson DN. The spatial and mechanical challenges of female meiosis. Mol Reprod Dev. 2011;78(10-11):769–77.

5. De Santis L, Cino I, Rabellotti E, Calzi F, Persico P, Borini A, et al. Polar body morphology and spindle imaging as predictors of oocyte quality. Reprod Biomed Online. 2005;11(1):36–42.

6. Rienzi L, Ubaldi F, Martinez F, Iacobelli M, Minasi MG, Ferrero S, et al. Relationship between meiotic spindle location with regard to the polar body position and oocyte developmental potential after ICSI. Hum Reprod. 2003;18(6):1289–93.

7. Chuang CH, Schlientz AJ, Yang J, Bowerman B. Microtubule assembly and pole coalescence: early steps in Caenorhabditiselegans oocyte meiosis I spindle assembly. Biol Open. 2020;9(6).

8. Speliotes EK, Uren A, Vaux D, Horvitz HR. The survivin-like C. elegans BIR-1 protein acts with the Aurora-like kinase AIR-2 to affect chromosomes and the spindle midzone. Mol Cell. 2000;6(2):211–23.

9. Romano A, Guse A, Krascenicova I, Schnabel H, Schnabel R, Glotzer M. CSC-1: a subunit of the Aurora B kinase complex that binds to the survivin-like protein BIR-1 and the incenp-like protein ICP-1. J Cell Biol. 2003;161(2):229–36.

10. Gong T, McNally KL, Konanoor S, Peraza A, Bailey C, Redemann S, et al. Roles of Tubulin Concentration during Prometaphase and Ran-GTP during Anaphase of C. elegans meiosis. bioRxiv. 2024.

11. Wu T, Dong J, Fu J, Kuang Y, Chen B, Gu H, et al. The mechanism of acentrosomal spindle assembly in human oocytes. Science. 2022;378(6621):eabq7361.

12. Harasimov K, Uraji J, Monnich EU, Holubcova Z, Elder K, Blayney M, et al. Actin-driven chromosome clustering facilitates fast and complete chromosome capture in mammalian oocytes. Nat Cell Biol. 2023;25(3):439–52.

13. Mitchison TJ, Maddox P, Gaetz J, Groen A, Shirasu M, Desai A, et al. Roles of polymerization dynamics, opposed motors, and a tensile element in governing the length of Xenopus extract meiotic spindles. Mol Biol Cell. 2005;16(6):3064–76.

14. Mossadeq LE, Bellutti L, Borgne RL, Canman JC, Pintard L, Verbavatz JM, et al. An interkinetic envelope surrounds chromosomes between meiosis I and II in C. elegans oocytes. bioRxiv. 2024.

15. Dumont J, Desai A. Acentrosomal spindle assembly and chromosome segregation during oocyte meiosis. Trends Cell Biol. 2012;22(5):241–9.

16. Lantzsch I, Yu CH, Chen YZ, Zimyanin V, Yazdkhasti H, Lindow N, et al. Microtubule reorganization during female meiosis in C. elegans. Elife. 2021;10.

17. Redemann S, Lantzsch I, Lindow N, Prohaska S, Srayko M, Muller-Reichert T. A Switch in Microtubule Orientation during C. elegans Meiosis. Curr Biol. 2018;28(18):2991–7 e2.

18. So C, Seres KB, Steyer AM, Monnich E, Clift D, Pejkovska A, et al. A liquid-like spindle domain promotes acentrosomal spindle assembly in mammalian oocytes. Science. 2019;364(6447).

19. Fabritius AS, Ellefson ML, McNally FJ. Nuclear and spindle positioning during oocyte meiosis. Curr Opin Cell Biol. 2011;23(1):78–84.

20. Yuan F, Alimohamadi H, Bakka B, Trementozzi AN, Day KJ, Fawzi NL, et al. Membrane bending by protein phase separation. Proc Natl Acad Sci U S A. 2021;118(11).

21. Wang Y, Li S, Mokbel M, May AI, Liang Z, Zeng Y, et al. Biomolecular condensates mediate bending and scission of endosome membranes. Nature. 2024;634(8036):1204–10.

22. Cavin-Meza G, Mullen TJ, Czajkowski ER, Wolff ID, Divekar NS, Finkle JD, et al. ZYG-9ch-TOG promotes the stability of acentrosomal poles via regulation of spindle microtubules in C. elegans oocyte meiosis. PLoS Genet. 2022;18(11):e1010489.

23. Harvey AM, Chuang CH, Sumiyoshi E, Bowerman B. C. elegans XMAP215/ZYG-9 and TACC/TAC-1 act at multiple times during oocyte meiotic spindle assembly and promote both spindle pole coalescence and stability. PLoS Genet. 2023;19(1):e1010363.

24. Marques A, Pedrosa-Harand A. Holocentromere identity: from the typical mitotic linear structure to the great plasticity of meiotic holocentromeres. Chromosoma. 2016;125(4):669–81.

25. Pitayu-Nugroho L, Aubry M, Laband K, Geoffroy H, Ganeswaran T, Primadhanty A, et al. Kinetochore component function in C. elegans oocytes revealed by 4D tracking of holocentric chromosomes. Nat Commun. 2023;14(1):4032.

26. Akiyoshi B, Sarangapani KK, Powers AF, Nelson CR, Reichow SL, Arellano-Santoyo H, et al. Tension directly stabilizes reconstituted kinetochore-microtubule attachments. Nature. 2010;468(7323):576–9.

27. Miller MP, Asbury CL, Biggins S. A TOG Protein Confers Tension Sensitivity to Kinetochore-Microtubule Attachments. Cell. 2016;165(6):1428–39.

28. Miller MP, Evans RK, Zelter A, Geyer EA, MacCoss MJ, Rice LM, et al. Kinetochore-associated Stu2 promotes chromosome biorientation in vivo. PLoS Genet. 2019;15(10):e1008423.

29. Zahm JA, Stewart MG, Carrier JS, Harrison SC, Miller MP. Structural basis of Stu2 recruitment to yeast kinetochores. Elife. 2021;10.

30. Desai A, Rybina S, Muller-Reichert T, Shevchenko A, Shevchenko A, Hyman A, et al. KNL-1 directs assembly of the microtubule-binding interface of the kinetochore in C. elegans. Genes Dev. 2003;17(19):2421–35.

31. Chen L, Maristany MJ, Farr SE, Luo J, Gibson BA, Doolittle LK, et al. Nucleosome Spacing Can Fine-Tune Higher Order Chromatin Assembly. bioRxiv. 2024.

32. Gibson BA, Blaukopf C, Lou T, Chen L, Doolittle LK, Finkelstein I, et al. In diverse conditions, intrinsic chromatin condensates have liquid-like material properties. Proc Natl Acad Sci U S A. 2023;120(18):e2218085120.

33. Cisneros-Soberanis F, Simpson EL, Beckett AJ, Pucekova N, Corless S, Kochanova NY, et al. Near millimolar concentration of nucleosomes in mitotic chromosomes from late prometaphase into anaphase. J Cell Biol. 2024;223(11).

34. Zhou H, Huertas J, Maristany MJ, Russell K, Hwang JH, Yao RW, et al. Multiscale structure of chromatin condensates explains phase separation and material properties. Science. 2025;390(6777):eadv6588.

35. Brouhard GJ, Stear JH, Noetzel TL, Al-Bassam J, Kinoshita K, Harrison SC, et al. XMAP215 is a processive microtubule polymerase. Cell. 2008;132(1):79–88.

36. Geyer EA, Miller MP, Brautigam CA, Biggins S, Rice LM. Design principles of a microtubule polymerase. Elife. 2018;7.

37. Bellanger JM, Gonczy P. TAC-1 and ZYG-9 form a complex that promotes microtubule assembly in C. elegans embryos. Curr Biol. 2003;13(17):1488–98.

38. Herman JA, Miller MP, Biggins S. chTOG is a conserved mitotic error correction factor. Elife. 2020;9.

39. Sathyapriya R, Vishveshwara S. Interaction of DNA with clusters of amino acids in proteins. Nucleic Acids Res. 2004;32(14):4109–18.

40. Bartas M, Cerven J, Guziurova S, Slychko K, Pecinka P. Amino Acid Composition in Various Types of Nucleic Acid-Binding Proteins. Int J Mol Sci. 2021;22(2).

41. Matthews LR, Carter P, Thierry-Mieg D, Kemphues K. ZYG-9, a Caenorhabditis elegans protein required for microtubule organization and function, is a component of meiotic and mitotic spindle poles. J Cell Biol. 1998;141(5):1159–68.

42. Woodruff JB, Ferreira Gomes B, Widlund PO, Mahamid J, Honigmann A, Hyman AA. The Centrosome Is a Selective Condensate that Nucleates Microtubules by Concentrating Tubulin. Cell. 2017;169(6):1066–77 e10.

43. Tilney LG, Porter KR. Studies on the microtubules in heliozoa. II. The effect of low temperature on these structures in the formation and maintenance of the axopodia. J Cell Biol. 1967;34(1):327–43.

44. Pallotto LM, Dilks CM, Park YJ, Smit RB, Lu BT, Gopalakrishnan C, et al. Interactions of Caenorhabditis elegans beta-tubulins with the microtubule inhibitor and anthelmintic drug albendazole. Genetics. 2022;221(4).

45. Balczon R, Prasain N, Ochoa C, Prater J, Zhu B, Alexeyev M, et al. Pseudomonas aeruginosa exotoxin Y-mediated tau hyperphosphorylation impairs microtubule assembly in pulmonary microvascular endothelial cells. PLoS One. 2013;8(9):e74343.

46. Currie JD, Stewman S, Schimizzi G, Slep KC, Ma A, Rogers SL. The microtubule lattice and plus-end association of Drosophila Mini spindles is spatially regulated to fine-tune microtubule dynamics. Mol Biol Cell. 2011;22(22):4343–61.

47. Kronja I, Kruljac-Letunic A, Caudron-Herger M, Bieling P, Karsenti E. XMAP215-EB1 interaction is required for proper spindle assembly and chromosome segregation in Xenopus egg extract. Mol Biol Cell. 2009;20(11):2684–96.

48. Sun M, Heald R. A DNA Crosslinker Collects Mitotic Chromosomes. Dev Cell. 2017;42(5):440–2.

49. Cuylen S, Blaukopf C, Politi AZ, Muller-Reichert T, Neumann B, Poser I, et al. Ki-67 acts as a biological surfactant to disperse mitotic chromosomes. Nature. 2016;535(7611):308–12.

50. Cuylen-Haering S, Petrovic M, Hernandez-Armendariz A, Schneider MWG, Samwer M, Blaukopf C, et al. Chromosome clustering by Ki-67 excludes cytoplasm during nuclear assembly. Nature. 2020;587(7833):285–90.

51. Hernandez-Armendariz A, Sorichetti V, Hayashi Y, Koskova Z, Brunner A, Ellenberg J, et al. A liquid-like coat mediates chromosome clustering during mitotic exit. Mol Cell. 2024;84(17):3254–70 e9.

52. Stamatiou K, Vagnarelli P. Chromosome clustering in mitosis by the nuclear protein Ki-67. Biochem Soc Trans. 2021;49(6):2767–76.

53. Remnant L, Kochanova NY, Reid C, Cisneros-Soberanis F, Earnshaw WC. The intrinsically disorderly story of Ki-67. Open Biol. 2021;11(8):210120.

54. Sobecki M, Mrouj K, Colinge J, Gerbe F, Jay P, Krasinska L, et al. Cell-Cycle Regulation Accounts for Variability in Ki-67 Expression Levels. Cancer Res. 2017;77(10):2722–34.

55. Takagi M, Ono T, Natsume T, Sakamoto C, Nakao M, Saitoh N, et al. Ki-67 and condensins support the integrity of mitotic chromosomes through distinct mechanisms. J Cell Sci. 2018;131(6).

56. Cidado J, Wong HY, Rosen DM, Cimino-Mathews A, Garay JP, Fessler AG, et al. Ki-67 is required for maintenance of cancer stem cells but not cell proliferation. Oncotarget. 2016;7(5):6281–93.

57. Samwer M, Schneider MWG, Hoefler R, Schmalhorst PS, Jude JG, Zuber J, et al. DNA Cross-Bridging Shapes a Single Nucleus from a Set of Mitotic Chromosomes. Cell. 2017;170(5):956–72 e23.

58. Margalit A, Brachner A, Gotzmann J, Foisner R, Gruenbaum Y. Barrier-to-autointegration factor--a BAFfling little protein. Trends Cell Biol. 2007;17(4):202–8.

59. Gorjanacz M, Klerkx EP, Galy V, Santarella R, Lopez-Iglesias C, Askjaer P, et al. Caenorhabditis elegans BAF-1 and its kinase VRK-1 participate directly in post-mitotic nuclear envelope assembly. EMBO J. 2007;26(1):132–43.

60. El Mossadeq L, Bellutti L, Le Borgne R, Canman JC, Pintard L, Verbavatz JM, et al. An interkinetic envelope surrounds chromosomes between meiosis I and II in C. elegans oocytes. J Cell Biol. 2025;224(3).

61. Lemaitre RP, Bogdanova A, Borgonovo B, Woodruff JB, Drechsel DN. FlexiBAC: a versatile, open-source baculovirus vector system for protein expression, secretion, and proteolytic processing. BMC Biotechnol. 2019;19(1):20.

62. Laband K, Lacroix B, Edwards F, Canman JC, Dumont J. Live imaging of C. elegans oocytes and early embryos. Methods Cell Biol. 2018;145:217–36.

63. Rios MU, Bagnucka MA, Ryder BD, Ferreira Gomes B, Familiari NE, Yaguchi K, et al. Multivalent coiled-coil interactions enable full-scale centrosome assembly and strength. J Cell Biol. 2024;223(4).

64. Gell C, Friel CT, Borgonovo B, Drechsel DN, Hyman AA, Howard J. Purification of tubulin from porcine brain. Methods Mol Biol. 2011;777:15–28.

